# Myosin in autoinhibited *off* state(s), stabilized by mavacamten, can be recruited via inotropic effectors

**DOI:** 10.1101/2023.04.10.536292

**Authors:** Weikang Ma, Carlos L. del Rio, Lin Qi, Momcilo Prodanovic, Srboljub Mijailovich, Christopher Zambataro, Henry Gong, Rafael Shimkunas, Sampath Gollapudi, Suman Nag, Thomas C. Irving

**Author notes:** authors contributed equally (Weikang Ma, Carlos L. del Rio). Corresponding authors: Suman Nag,; Carlos L. del Rio,; Weikang Ma.

## Abstract

Mavacamten is a novel, FDA-approved, small molecule therapeutic designed to regulate cardiac function by selectively but reversibly inhibiting the enzymatic activity of myosin. It shifts myosin towards ordered *off* states close to the thick filament backbone. It remains unresolved whether mavacamten permanently sequesters these myosin heads in the *off* state(s) or whether these heads can be recruited in response to physiological stimuli when required to boost cardiac output. We show that cardiac myosins stabilized in these *off* state(s) by mavacamten are recruitable by Ca^2+^, increased heart rate, stretch, and β-adrenergic (β-AR) stimulation, all known physiological inotropic effectors. At the molecular level, we show that, in presence of mavacamten, Ca^2+^ increases myosin ATPase activity by shifting myosin heads from the reserve super-relaxed (SRX) state to the active disordered relaxed (DRX) state. At the myofilament level, both Ca^2+^ and passive lengthening can shift ordered *off* myosin heads from positions close to the thick filament backbone to disordered *on* states closer to the thin filaments in the presence of mavacamten. In isolated rat cardiomyocytes, increased stimulation rates enhanced shortening fraction in mavacamten-treated cells. This observation was confirmed *in vivo* in telemetered rats, where left-ventricular dP/dt_max,_ an index of inotropy, increased with heart rate in mavacamten treated animals. Finally, we show that β-AR stimulation *in vivo* increases left-ventricular function and stroke volume in the setting of mavacamten. Our data demonstrate that the mavacamten-promoted *off* states of myosin in the thick filament are activable, at least partially, thus leading to preservation of cardiac reserve mechanisms.

**Significance statement:** Mavacamten is the first myosin-targeted small molecule inhibitor approved by the FDA to treat obstructive hypertrophic cardiomyopathy by attenuating myocardial hyperdynamic contraction. The recruitment of cardiac contractility is, however, vital to ensure sufficient cardiac output during increased physiological demand. Here we show that major inotropic effectors are at least partially preserved in the setting of mavacamten, resulting in maintenance of cardiac reserve mechanisms. These results not only suggest an alternative mechanistic explanation, beyond mere LV outflow tract obstruction removal, for the clinically observed increase in peak oxygen uptake with exercise in HCM patients receiving mavacamten, but also lay the groundwork for a potential methodology to investigate the sarcomeric basis of chronotropic incompetence in disease states to motivate new therapeutic interventions.

## Introduction

The heart is the powerhouse of the circulatory system, and its stroke volume is precisely regulated by various inotropic effectors to fulfill the body’s demands. Four foremost positive inotropic effectors exist in the heart: 1) increase in calcium (Ca^2+^) concentration via the action of L-type Ca^2+^ channels (1); 2) via increased heart rate or frequency, known as the Bowditch effect or the Treppe phenomenon (2, 3); 3) increased diastolic ventricular preload, also known as length-dependent activation (LDA)(4); and 4) β-adrenergic stimulation that increases cyclic AMP, activating protein kinase A (PKA) which subsequently phosphorylates regulatory proteins in the heart muscle resulting in increased cardiac contractility and, consequently, cardiac output (5). The ability to maintain these positive inotropic effects is vital for the heart to function properly in response to increased physiological demands, and deficits in the heart’s response to both inotropes and increased demand is a common feature of both cardiomyopathies (6) and/or acquired cardiovascular diseases (7).

Such is the case in hypertrophic cardiomyopathy (HCM), a myocardial disorder characterized by left ventricular (LV) hypertrophy, hyperdynamic contraction, and impaired relaxation (8). The cardiac changes in HCM often manifest as hindered exercise capacity, as reflected by reduced peak oxygen uptake (pVO_2_) and cardiac output recruitment. The root causes of the disease in a significant fraction of HCM patients lie within the sarcomere, the basic contractile apparatus of the heart. Approximately 45% of affected individuals, and a significant portion of those with a family history of clinical disease, have a mutation(s) in one or more genes encoding sarcomeric proteins (8). It has been shown that these sarcomeric mutations lead to excessive actin-myosin interactions, or cross-bridges, arising from a pathogenic shift of myosin heads from reserve and energy-sparing *off* states towards *on* states (9, 10). Other mutations outside of the sarcomere may also contribute to HCM (11).

Mavacamten (commercially known as Camzyos) is an FDA approved small molecule developed to address this pathogenicity by selectively attenuating the ATPase activity of cardiac myosin(12). This targeted myosin inhibition results in a novel pharmacological profile that is capable of normalizing contraction and relaxation properties, as demonstrated in both preclinical and phase-II and phase-III clinical studies, being approved, now in multiple countries for the treatment of patients with obstructive HCM (13-15). A comprehensive review on the mechanism of mavacamten has been recently published (16). More importantly, despite reducing the intrinsic activity of myosin, the sarcomere motor of the heart, mavacamten has been shown to increase pVO_2_ in patients with obstructive HCM (14, 17) and in a subset of non-obstructive HCM patients (18). Thus, as pVO_2_ reflects the ability to recruit forward flow and stroke volume during exercise, the mechanistic basis for these effects in the setting of an agent designed to attenuate cardiac myosin function, remains to be elucidated and may lie within the molecular impacts of mavacamten in the sarcomere.

Although the atomic structure of myosin with mavacamten bound has yet to be published, electron microscopy (13) and SAXS (19) studies show that mavacamten stabilizes a compact folded-back structure of myosin (13) that may not be exactly similar to the interacting-heads motif (IHM) configuration of myosin that is seen in relaxed skeletal, cardiac, and smooth muscle myosin from diverse organisms (16, 20). Additionally, mavacamten stabilizes the biochemically energy-sparing auto-inhibited super-relaxed (SRX) state(s) of myosin (13, 19, 21, 22). Consistent with these *in vitro* observations, *ex vivo* low-angle X-ray diffraction studies of porcine cardiac muscle fibers after mavacamten treatment show more heads sequestered closer to the thick filament, thereby increasing the quasi-helical ordering of myosin heads along the thick filament backbone (13, 23, 24). Whether myosin in the SRX state(s) is necessarily in the canonical IHM structural state (10, 20, 25), or whether structurally-defined sequestered heads are necessarily in the biochemically defined SRX state or necessarily adopt the IHM configuration remain open questions in the field (see(26)). Regardless, this body of evidence suggests that mavacamten induces a population of myosin heads to adopt sequestered *off* state(s) with low probability (availability) to participate in contraction under normal physiological conditions. This raises the question, however, whether these mavacamten-stabilized *off* state(s) can be recruited during increased workload in order to increase forward flow and meet the physiological demands of mavacamten-treated patients under physical stress.

Some hints of this potential recruitability can be found in the literature (12, 27), where mavacamten has been shown to preserve, at least partially, responsiveness to exercise and β-AR stimulation in *vivo*, as well as length-dependent activation (LDA) in *vitro*(23, 28, 29). LDA is a key feature of cardiac muscle constituting the cellular basis of the Frank-Starling mechanism, wherein the muscle responds with increased Ca^2+^ sensitivity and force to increases in length (preload), whereby the heart increases its cardiac output (4). Functionally, mavacamten is shown to preserve, at least partially, LDA (23, 28, 29), suggesting that the myosin heads sequestered in the *off* SRX state(s) induced by mavacamten are potentially available for recruitment. To test the recruitability of mavacamten-treated myocardium, we designed experiments ranging from molecular level, studies on cardiac myosin synthetic thick filaments, cardiac myofibrils using SRX/DRX and ATPase assays to sarcomere level studies on permeabilized porcine myocardium using small-angle X-ray diffraction, computational simulations, and in *vivo* experiments in conscious and anesthetized animals. Collectively, these studies demonstrate that the myosin sequestered *off* states stabilized by mavacamten are controllable and can be recruited, at least partially, by 1) direct increase in Ca^2+^ in *vitro* or 2) via increase in Ca^2+^ due to increased heart rate in *vivo*, 3) sarcomere stretch as well as 4) β-AR stimulation.

## Methods

### β**-cardiac full-length myosin protein purification**

β-cardiac full-length myosin from the bovine left ventricle is isolated following established methods described previously (30). Following this, full-length myosin is dialyzed in a buffer containing 10 mM PIPES (pH 6.8), 300 mM KCl, 0.5 mM MgCl_2_, 0.5 mM EGTA, 1 mM NaHCO_3_, and 1 mM DTT and stored at −80°C. Based on densitometry analysis of SDS-PAGE, the purity of the myosin preparation varies between 90 and 95%, with negligible actin contamination. Also, the basal myosin ATPase from these preparations always turns out to be 0.03 ± 0.01 s^−1^, suggesting negligible actin-activation. Bovine cardiac myofibrils were prepared following the methods described previously(31) and were stored at −80°C in a buffer containing 10 mM PIPES (pH 6.8), 30 mM KCl, 0.5 mM MgCl_2_, 0.5 mM EGTA, 1 mM NaHCO_3_, and 1 mM DTT.

#### Reconstitution of synthetic myosin thick filaments

Full-length myosin remains fully soluble in a buffer of high ionic strength (300 mM). However, many previous studies had previously shown that the full-length myosin could spontaneously self-assemble into bipolar thick filaments when the ionic strength of the buffer was reduced to 150 mM or below (32, 33). This method was used to construct thick filaments with some modifications in this study. Briefly, the ionic strength of the myosin sample and the myosin concentration were first adjusted to the desired value by diluting it with a buffer containing 20-mM Tris-HCl (pH 7.4), 0-mM KCl, 1-mM EGTA, 3-mM MgCl2, and 1-mM DTT. After dilution, the myosin sample was incubated for 2 h on ice to form thick filaments before using it for experiments. In this study, such reconstituted myosin filaments will be referred to as STFs. Alternatively, STFs were also made by slowly dialyzing full-length myosin into a low ionic (150 mM or below) strength buffer. Quantitatively, the measurements reported in STFs were not different between these two methods.

#### Steady-state ATPase measurements

Measurements of basal myosin and myofibrillar ATPase activity as a function of Ca^2+^ were performed at 23 °C on a plate-based reader (SpectraMax 96- well) using an enzymatically coupled assay as described previously (34). The buffer conditions used were 12 mM Pipes (pH 6.8), 2 mM MgCl_2_, 10 mM KCl, and 1 mM DTT. For the basal ATPase experiments with STFs, a final myosin concentration of 1 µM was used. When using myofibrils, approximately 40% of the total weight was assumed to be from myosin, and accordingly, the myofibril amount was loaded to attain a final myosin concentration of 1 μM. A concentrated stock (20 mM) of mavacamten and MYK-7660 (a thin filament inhibitor) was first prepared using dimethyl sulfoxide (DMSO), and this was used to attain the desired concentration in the final buffer samples. A final DMSO concentration of 2% was achieved in all samples. Data sets recorded by the instrument-compatible software package, SoftMax Pro, were exported to GraphPad Prism and analyzed.

#### Biochemical SRX/DRX measurements in reconstituted myosin synthetic thick filaments

β- cardiac full-length myosin from the bovine left ventricle was isolated following established methods described elsewhere(30). Bipolar synthetic thick filaments (STF) from this full-length myosin were prepared (∼ 0.4 μM) as described previously (22). Briefly, full-length bovine β- cardiac myosin remains fully soluble in a buffer of high ionic strength (300 mM) but spontaneously self-assembles into bipolar thick filaments at lower ionic strengths (22). For each experiment, 10 µM of full-length myosin in 300 mM KCl buffer was diluted to 1 µM in KCl buffer containing 20 mM Tris-HCl (pH 7.4), 0 mM KCl, 1 mM EGTA, 3 mM MgCl2, and 1 mM DTT, to achieve a final KCL concentration of 30 mM. To allow for thick filament formation, the sample was incubated for 2 h on ice, and then used in the experiments.

Single ATP turnover kinetic experiments using a fluorescent 2′/3′-O-(N-Methylanthraniloyl) (mant)-ATP were conducted in a 96-well plate fluorescence plate reader as described earlier (22). This assay measures fluorescent nucleotide release rates after incubation of myosin preparations with mant-ATP and chased with excess unlabeled ATP. In the first step, 100 μl of 0.8-μM myosin was combined with 50 μl of 3.2-μM mant-ATP in a UV-transparent fluorescence plate and the reaction was aged for 60 s to allow binding and hydrolysis of mant-ATP to inorganic phosphate and mant-ADP. In the second step, mant-nucleotides were chased with nonfluorescent ATP by adding 50 μl of 16-mM nonfluorescent ATP to the above mixture, and the resulting fluorescence decay due to mant-nucleotide dissociation from myosin was monitored over time. The composition of the final buffer was 20-mM Tris-HCl (pH 7.4), 30-mM KCl, 1- mM EGTA, 3-mM MgCl2, and 1-mM DTT. The concentrations of myosin, mant-ATP, and nonfluorescent ATP attained in the final mixture were 0.4 μM, 0.8 μM, and 4 mM, respectively. The final concentration of DMSO was 2% in all samples. The excitation wavelength for monitoring the mant-nucleotides was 385 nm. The emission was acquired using a long-pass filter with a cutoff at 450 nm. Data were acquired at a frequency of 4.25 Hz for the first 500 points, followed by 1 Hz for the next 430 points and finally at 0.16 Hz for the rest of the duration. All experiments were performed at 25 °C. The fluorescence decay profiles, obtained during the chase phase characteristically depict two phases, a fast phase followed by a slow phase. This data was fit to a bi-exponential function to estimate four parameters corresponding to fast and slow phases—A_fast_, k_fast_, A_slow_, and k_slow_—where A represents the normalized % amplitude and k represents the observed ATP turnover rate of each phase. The fast and the slow phases correspond to the myosin activity in the DRX and SRX states, respectively.

#### X-ray diffraction measurements on permeabilized porcine myocardium

Isolated hearts (N =2) from male Yucatan pig were provided by Exemplar Genetics Inc. Humane euthanasia and tissue collection procedures were approved by the Institutional Animal Care and Use Committees at Exemplar Genetics. The procedures were conducted according to the principles in the "Guide for the Care and Use of Laboratory Animals", Institute of Laboratory Animals Resources, Eighth Edition (35), the Animal Welfare Act as amended, and with accepted American Veterinary Medical Association (AVMA) guidelines at the time of the experiments(36). Left ventricular myocardium samples were prepared as described previously (37, 38). Briefly, samples of frozen left ventricular myocardium were permeabilized in relaxing solution (2.25 mM Na_2_ATP, 3.56 mM MgCl_2_, 7 mM EGTA, 15 mM sodium phosphocreatine, 91.2 mM potassium propionate, 20 mM Imidazole, 0.165 mM CaCl_2_, creatine phosphate kinase 15 U/ml) containing 15 mM 2,3-Butanedione 2-monoxime (BDM) and 1% Triton-X100 and 3% dextran at pH 7) at room temperature for 2-3h). Muscles were then washed with fresh relaxing solution for ∼ 10 minutes, repeated 3 times. The tissue was dissected into ∼200 *μm* diameter fiber bundles and clipped with aluminum T-clips. The preparations were then stored in cold (4°C) relaxing solution with 3% dextran for the day’s experiments. X-ray diffraction experiments were performed at the BioCAT beamline 18ID at the Advanced Photon Source, Argonne National Laboratory (39). The muscles were incubated in a customized chamber, and experiments were performed at 28-30 °C. For the results presented in Fig. 2, the muscle was held at a sarcomere length of 2.3 µm and the X-ray patterns were collected at seven different Ca^2+^ concentrations (pCa 8, pCa 6.4, pCa 6, pCa 5.8, pCa 5.6, pCa 5.3, and pCa 4.5) in the presence of 100 μM thin filament inhibitor (MYK-7660), and 2 μM of mavacamten (in the mavacamten group) to compare to the results from previous studies in the absence of these compounds (24).

We used the thin filament inhibitor, MYK-7660, in this study simply as a tool to decouple the Ca^2+^ effects mediated via the thin and the thick filament. The exact mechanism of inhibition remains, however, beyond the scope of this study. A detailed description of the behavior of this molecule was recently published (24). In our previous study, we showed that MYK-7660 did not affect actin-myosin activity unless troponin and tropomyosin complexes are present. This result indicates that MYK-7660 does not bind to myosin. 100 μM of MYK-7660 was used to saturate and block all Ca^2+^-mediated activity of the thin filament so that any Ca^2+^- mediated activity observed from the system could be attributed to its effect on the thick filaments. For the results presented in Fig. 4, two groups of X-ray patterns were collected at a sarcomere length of 2.1 µm and then at a sarcomere length of 2.3 µm in the presence and absence of 50 μM mavacamten respectively. X-ray patterns were collected on a MarCCD 165 detector (Rayonix Inc., Evanston IL) with a 1 s exposure time. 2-3 patterns were collected under each condition. Reflection spacings and intensities were extracted from these patterns using data reduction programs belonging to the open-source MuscleX software package developed at BioCAT (40) and averaged. Briefly, the equatorial reflections were measured by the "Equator" routine in MuscleX as described previously (41). Briefly, the equatorial intensity trace was reduced to a one-dimensional projection by drawing a box along the equator. After removing the diffused background, the intensity distribution along the equator was fit using a Marquardt– Levenberg algorithm to calculate the peak intensities. The intensities (23) and spacings (42) of meridional and layer line reflections were measured by the "Projection Traces" routine in MuscleX. Briefly, the meridional reflections are reduced into one-dimensional intensity profiles by drawing a box along the meridian. The target peak positions were estimated as the centroid of the intensity of the top half of the diffraction peak as described previously (43). The intensity of the target reflections were modeled as a single Gaussian peak.

#### Isolated Myocytes (Bowditch)

Rat experiments were done in compliance with protocols approved by the Institutional Animal Care and Use Committees at Bristol-Myers Squibb (formerly MyoKardia Inc.). Primary left-ventricular cardiomyocytes were isolated were from adult Sprague Dawley rats via enzymatic digestion in a Langendorff apparatus as previously described (12). Briefly, the rat heart was removed and cannulated on a perfusion system with Tyrode’s buffer in the presence of collagenase. Subsequently, aliquots of cardiomyocyte suspensions in Tyrode’s buffer solution kept at room temperature (20-22 °C) were placed into the perfusion chamber on an inverted microscope, and cardiomyocytes were allowed to settle and adhere to the chamber glass coverslip for 5 minutes. Tyrode’s solution containing DMSO (control, with 0.05% DMSO) was then superfused into the chamber at 1.5 mL/min and solution temperature gradually increased to and kept at 31-32 °C by in-line pre-heater temperature controller (Cell MicroControls, Norfolk, VA, USA). All measurements were performed at 31-32 °C. Once at set temperature, cardiomyocyte pacing was started by electrical stimulations between two platinum electrodes at 1 Hz and 6 V for 2 ms (MyoPacer, IonOptix, Westwood, MA, USA). Cardiomyocytes were paced for at least 5 minutes to reach steady state contraction prior to experiments. Solutions containing either DMSO or mavacamten (0.1 µM) were superfused into the chamber for 5 minutes followed by progressive increases in pacing rate (to 2 Hz, 3 Hz, and 4 Hz). Measurements were taken from the last 30 seconds of each pacing/exposure period. Concurrent measurements of cell-length and cytosolic calcium fluorescence were acquired from single cardiomyocytes for each experiment recording.

For calcium fluorescence measurements, cardiomyocytes were incubated with 1 µM Fura-2- acetoxymethyl ester (Fura-2 AM, cell permeant, ThermoFisher) and 0.075% Pluronic F-127 (20% in DMSO, ThermoFisher) for 15 minutes at room temperature. Cardiomyocytes were then washed with fresh Tyrode’s solution and incubated for 15 minutes at room temperature to allow for de-esterification. Calcium fluorescence was measured by excitation of the Fura-2 fluorophore by separate light-emitting diode (LED) light sources producing light in the UV spectra at 340 and 380 nm. Bandpass filters on each LED light restricted excitation light at 340 and 380 nm. Digital modulation generated alternating excitation wavelengths at 250 Hz, and the light was directed into the perfusion chamber. The framing aperture was centered around single cardiomyocytes to ensure that the fluorescence signal was acquired from only the selected cardiomyocyte. Emitted light at 510 nm passed through a dichroic mirror and directed to a photomultiplier tube (PMT) where the photon count was amplified and converted to a digital signal for fluorescence quantification. The output signal for the Fura-2 ratiometric method consisted of 250 ratios per second and recorded with IonWizard software (IonOptix). For contractility measurements, cardiomyocytes were imaged through a 40X objective in an inverted microscope. Cardiomyocyte contractility was measured by video-based CMOS sensor (MyoCam, IonOptix) quantification of changes in SL by fast Fourier transform (FFT) of a region on interest (ROI) selected on a longitudinal section of the cardiomyocyte with a clear striation pattern. The ROI was kept in the same location on the cardiomyocyte for the duration of the experiment. Video capture produced SL measurements at 500 Hz. SL signals were recorded with IonWizard software (IonOptix) concurrent with calcium fluorescence. For analysis, a segment of steady-state cardiomyocyte contraction was selected for each protocol period: 1) pre-treatment (PRE), 2) either DMSO (Ctrl), or mavacamten (MAVA) at 1Hz, and 3) for stimulations at 2, 3, and 4Hz. At least 10 calcium transients were included for each analyzed period.

#### *In vivo* studies (Bowditch)

The rat experiments were done in compliance with protocols approved by the Institutional Animal Care and Use Committees at Bristol-Myers Squibb (formerly MyoKardia Inc.). Conscious cardiovascular responses to acute myosin-modulation via mavacamten were studied in telemetered adult male Sprague Dawley rats instrumented with an implantable radio-telemetry unit providing both systemic arterial (abdominal aorta, AoP) and left-ventricular pressures (model HD-21; Data Sciences International, Inc.). These animals were (orally) administered either DMSO (Ctrl, 0 mg/kg, n = 11) or mavacamten (MAVA: 1.0 mg/kg PO, n = 6). In these experiments, LV hemodynamics were recorded continuously via telemetry in conscious free-roaming animals both prior to (up 1 hour) and following (up to 24 hours) each dosing. Heart rate (HR) as well as the peak rates of pressure rise (dP/dtmax) and decline (dP/dtmin) were derived/measured from LV pressure signals, and are reported for the post-dosing period.

#### *In vivo* studies (β-adrenergic stimulation)

The pig experiments were done in compliance with protocols approved by the Institutional Animal Care and Use Committees at Exemplar Genetics. Four healthy Yucatan male pigs (58.0±9.5 Kg), were chemically-restrained (Telazol/Xylazine/Isoflurane) and underwent transthoracic echocardiographic examinations. Left-ventricular end-systolic and end-diastolic dimensions were measured from 2D-guided M-mode images in the parasternal long-and mid-papillary short-axis views; volumes and left-ventricular ejection fraction (LVEF) were calculated from internal dimension via the Teichholz method. End-diastolic volumes were indexed to body-surface area (EDVI = EDV/BSA) as described before (44) to account for differences in body weight. These measurements were then repeated after mavacamten administration (0.23 ± 0.03 mg/kg IV, at ∼2mL/kg). These measurements were then repeated after dobutamine administration (+DOB, 5 µg/kg/min) for 5 min to evaluate functional recruitment.

#### Computational simulations of thick filament regulation by calcium

For simulations of thick filament regulation by calcium, we used a recently updated version of the MUSICO computational platform (45-47) with parameters modified from the previous studies to account for differences between rodent and human myocardium and porcine myocardium. The full set of parameters used in the model is given in Table S1. The model incorporated into MUSICO accounts for all known interactions between sarcomere proteins, e.g., myosin with actin, the troponin-tropomyosin complex with the actin surface, and proteins with ions and small molecules such as Ca^2+^ in the context of a spatially explicit, three-dimensional (3D) structure of the sarcomere (45-47). The kinetic schemes describing these interactions include two major parts: (1) calcium binding to troponin and the regulation of availability of actin sites for myosin binding; (2) kinetics of the crossbridge cycle including interactions of myosin heads with actin and with the thick filament backbone. The first process is defined by four states that describe the thin filament-based regulation that binding of calcium to cTnC to form a Ca-cTnC complex that turn on thin filament. The second process is defined by a six-state crossbridge cycle that includes five biochemical states consistent with observed structural states: M.D.Pi, A.M.D.Pi, A.M.D, A.M, M.T and an additional sixth state, the so-called "parked state" (PS), represents the interaction of myosin heads with the thick filament backbone where they are unable to interact with actin. The transition rate from the PS to M.D.Pi is assumed to be strongly dependent on [Ca^2+^], i.e. 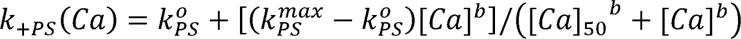 where 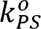 is baseline rate from parked state, 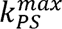 is the rate at high calcium, e.g. at pCa=4.5, b is Hill coefficient of the rate sigmoidal rise and[*Ca*]_50_ is calcium concentration when *k*_+*PS*_ is equal to 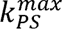/2 (45-47). In performing the simulations, the model incorporated known structural (lattice spacing, sarcomere length) and functional parameters (force-pCa curve). The simulations then varied the adjustable parameters (*k*_−*ps*_, 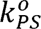, 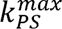, Hill coefficient, *b*, and reverse rate, *k*_−*ps*_) to provide good fits to the data in control experiments. With mavacamten, only *k*_−*ps*_ needed to be adjusted to quantitatively match the data. Since this is not a parametric model, there is no minimization process, so conventional goodness of fit statistics are not meaningful. See Supplemental Information (SI) for further details.

#### Statistics

Statistical analyses were performed using GraphPad Prism 9 (Graphpad Software). The relative changes versus pCa curves in Fig. 2 were fit to a four-parameter modified Hill equation (minimum response + (maximum response - minimum response)/(1+10^h^. (pCa50-pCa))) (48), where pCa_50_ is the calcium concentration yielding a response halfway between the minimum and maximum values reported in the article. Mann-Whitney unpaired nonparametric t-test were performed on the data in Fig. 2 between the thin filament inhibitor group and mavacamten groups. Wilcoxon matched-pairs nonparametric t-test were performed on the data in Fig. 3 within control and mavacamten groups. Mann-Whitney unpaired nonparametric t-test were performed on the data in Fig. 3 between control and mavacamten groups. The effects of mavacamten in isolated myocytes and in vivo were evaluated via a one-factor analysis of variance (ANOVA) with a repeated-measures design and set predetermined comparisons (e.g., DOSE (+mavacamten) vs. PRE (-mavacamten) and vs. DOSE (+mavacamten) +DOB). In addition, multiple linear regression analyses were used to compare the HR-dependent effects in dP/dt_max_ and dP/dt_min_. The significance level was set at 0.05. Symbols on figures: ns: not significant, *: p<0.05, **: p<0.01, ***: p<0.001,****: p<0.0001. N and n indicate biological and technical repeats respectively.

## Results

### Ca^2+^ destabilizes the SRX state(s) of myosin in thin-filament-free reconstituted synthetic myosin thick filaments

We have recently reported that Ca^2+^ binding to myosin filaments in reconstituted cardiac synthetic thick filaments (STF) can directly destabilize myosin SRX state(s) towards the DRX state(s) (24). Here we studied Ca^2+^-mediated destabilization in the presence of mavacamten, an agent capable of arresting myosin heads in the SRX state (Fig. 1a). The basal myosin SRX population in the STF system at pCa 10 is 10 ± 5 % (n=3), which progressively increases with increasing concentration of mavacamten. The IC_50_ of this transition is 1.2 ± 0.5 μM (Fig. 1a). Consistent with our earlier reports of a Ca^2+^-mediated destabilization mechanism(24), the myosin SRX population in the control sub-group progressively decreases with an increase in Ca^2+^ with a pCa_50_ of 5.5 (5.3 to 5.6 for 95% CI). Interestingly, this mechanism of Ca^2+^-mediated destabilization of the SRX population holds at all mavacamten concentrations tested (0.5, 1, and 2 μM) (Fig. 1b). These results suggest that while mavacamten puts more myosin into the SRX state(s) (Fig. 1b, left, upwards arrow), this effect is antagonized by increasing Ca^2+^ concentrations (Fig. 1b, right, upwards arrow). The pCa_50_ of the Ca^2+^-mediated destabilization in the presence of various concentrations of mavacamten lies in the range of 5.2 to 5.6 (4.9 to 5.7 for 95% CI) (n=3), suggesting that cytosolic Ca^2+^ concentrations traditionally associated with systole can destabilize mavacamten-induced SRX states. Taken together, these data signify that myosin heads are not permanently locked even at saturating doses of mavacamten (2 μM) but instead are recruitable as needed.

**Figure 1.**
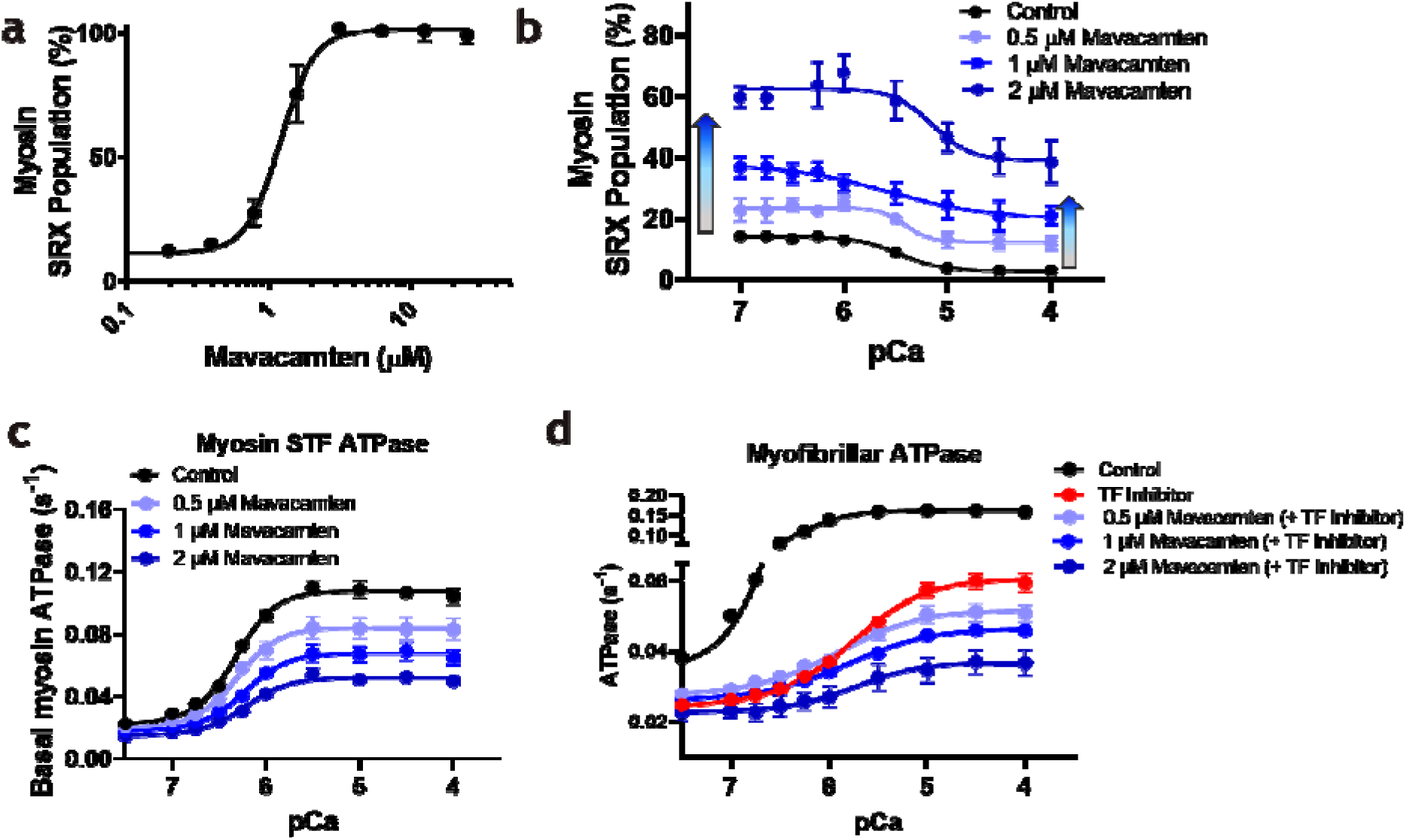
The biochemical SRX states of myosin stabilized by mavacamten are recruitable by Ca^2+^. **a**, Absolute percentage (%) of myosin SRX population in synthetic thick filaments (STF) reconstituted from bovine cardiac full-length myosin. The IC_50_ for this transition is 1.2 ± 0.5 μM. **b**, Absolute % myosin SRX population in STF in the absence (Control (2% DMSO), black) and the presence of increasing concentrations of mavacamten (0.5, 1 and 2 μM; increasing darker shades of blue) at different Ca^2+^ concentrations. Ca^2+^ destabilizes the myosin SRX state (s), otherwise stabilized by mavacamten. While mavacamten puts more myosins in the SRX state(s) at all concentrations (left, upwards arrow), this effect was blunted by increasing Ca^2+^ concentrations (right, upwards arrow). **c**, Absolute basal myosin ATPase in STF in the absence (Control (2% DMSO), black) and the presence of 0.5, 1, and 2 μM mavacamten concentrations (profiles in blue) at different Ca^2+^ concentrations. **d**, Absolute myosin ATPase in bovine cardiac myofibrils in the absence (Control (2% DMSO), black), and in the presence of 100 μM of thin filament inhibitor without (TF inhibitor, red) and with 0.5, 1, and 2 μM mavacamten concentrations (profiles in blue) at different Ca^2+^ concentrations.

Consistent with this mechanism, the basal ATPase activity of myosin in STF progressively increases with an increase in Ca^2+^ with a pCa_50_ of 6.32 (6.28 to 6.37 for 95% CI) (n=8) in the control sub-group (Fig. 1c). This mechanism of Ca^2+^-mediated activation of myosin ATPase also holds at 0.5, 1, and 2 µM mavacamten concentrations with pCa_50_ values of 6.31 (6.28 to 6.37 for 95% CI), 6.23 (6.18 to 6.28 for 95% CI), and 6.22 (6.14 to 6.30 for 95% CI) (Fig. 1c), respectively, suggesting that myosin heads inhibited by mavacamten can be re-activated with increasing Ca^2+^ concentrations. This premise is further tested in a more complex bovine cardiac myofibril system, encompassing both the thick and the thin filament components of the sarcomere. Increasing Ca^2+^ is expected to increase the myofibrillar ATPase via activating the thin-filament troponin-tropomyosin regulatory system (Fig. 1d, black; Control group; pCa_50_ of 6.36 and 6.32 to 6.40 for 95% CI). The Ca^2+^-dependent direct thick filament activation mechanism as demonstrated in the STF system above can be isolated from myofibrillar system by using a 100 µM thin filament inhibitor (MYK-7660)(24) to inhibit acto-myosin interactions presumably by stabilizing the Blocked or Closed states (49) of the thin filaments, thus abolishing active contraction. Our data shows that Ca^2+^-dependent thick filament activation, although blunted, was retained in the absence of active force (Fig. 1d, red; thin filament inhibitor; pCa_50_ of 5.78 and 5.73 to 5.83 for 95% CI), suggesting that a part of the Ca^2+^-mediated activation of the sarcomere lies in the thick filament, consistent with our previous report (24). Increasing mavacamten concentrations in the presence of 100 μM MYK-7660, in general, progressively blunted the ATPase activity due to myosin inhibition, but, very interestingly, was partially rescued by increasing Ca^2+^ (Fig. 1d, traces in blue). The pCa_50_ values of ATPase in myofibrils with 100 μM MYK-7660 treated with 0.5, 1, and 2 µM concentrations of mavacamten lies in the range of 5.76 to 5.93 (5.62 to 6.04 for 95% CI) (n=3). We note that it is also possible that at this high concentration of MYK-7660 and mavacamten, the sarcomere could be inhibited by other non-specific mechanisms, which would only lead to an underestimation of the degree of rescue by Ca^2+^, further bolstering our hypothesis.

### Ca^2+^ mediated myosin filament structural changes are preserved in the presence of mavacamten

We have previously shown that Ca^2+^ can directly move myosin heads from ordered *off* states to disordered *on* states in the absence of active force (24). Here we investigated how myosin thick filaments react to increasing Ca^2+^ concentration in the presence of mavacamten (blue curves in Fig. 2) and compared it to previously reported data (24) in the absence of mavacamten with (TF inhibitor, red curve in Fig. 2) and without thin filament inhibitor (Control, black curve in Fig. 2). The equatorial intensity ratio (I_1,1_/I_1,0_) is an indicator of the proximity of myosin heads to actin(26). The change in I_1,1_/I_1,0_ (Δ I_1,1_/I_1,0_) vs. pCa curve shows a sigmoidal shape (Fig. 2b) with a pCa_50_ of 5.7 for the Mavacamten group (N =2 and n = 7), the same as in the thin filament inhibitor and Control groups as previously reported (24). The increase in ΔI_1,1_/I_1,0_ with increasing Ca^2+^ concentration indicates that myosin heads move radially away from the thick filament backbone even in the presence of mavacamten. Relaxed myosin heads are quasi-helically ordered on the surface of the thick filament producing the myosin-based layer line reflections(26). The first-order myosin-based layer line (MLL1) arising from the helically ordering of myosin heads is disrupted when myosin heads are turned *on* to participate in contraction, leading to a decrease in the intensities of MLL1 (I_MLL1_). Similarly, the intensity of the third-order myosin-based meridional reflection (I_M3_) decreases when myosin heads become disordered (26, 50). I_M3_ (Fig. 2c) and I_MLL1_ (Fig. 2d) decays as a function of increasing Ca^2+^ concentration yields a pCa_50_ of 6.14 (5.93 to 6.38 for 95% CI) and 6.01 (5.84 to 6.20 for 95% CI) respectively in the presence of mavacamten. The spacing of the sixth-order meridional reflection (S_M6_) reports the periodicity of the thick filament backbone with increases in S_M6_ indicating extension of the thick filament backbone(42). The degree of thick filament lengthening has been shown to correlate with the degree of myosin heads being in the *on* states(51) and increases with increasing Ca^2+^ concentration with a pCa_50_ of 6.35 (>5.97 for 95% CI) for the mavacamten groups (Fig. 2e). In response to increasing Ca^2+^ concentration, the changes of I_1,1_/I_1,0_ (Fig. 2b) and S_M6_ (Fig. 2e) are larger in the control group while the changes of I_M3_ (Fig. 2c) and I_MLL1_ (Fig. 2d) are comparable among groups because I_1,1_/I_1,0_ and S_M6_ are sensitive to the number of active cross-bridges binding to actin while I_M3_ and I_MLL1_ are not necessarily coupled with active force.

**Figure 2.**
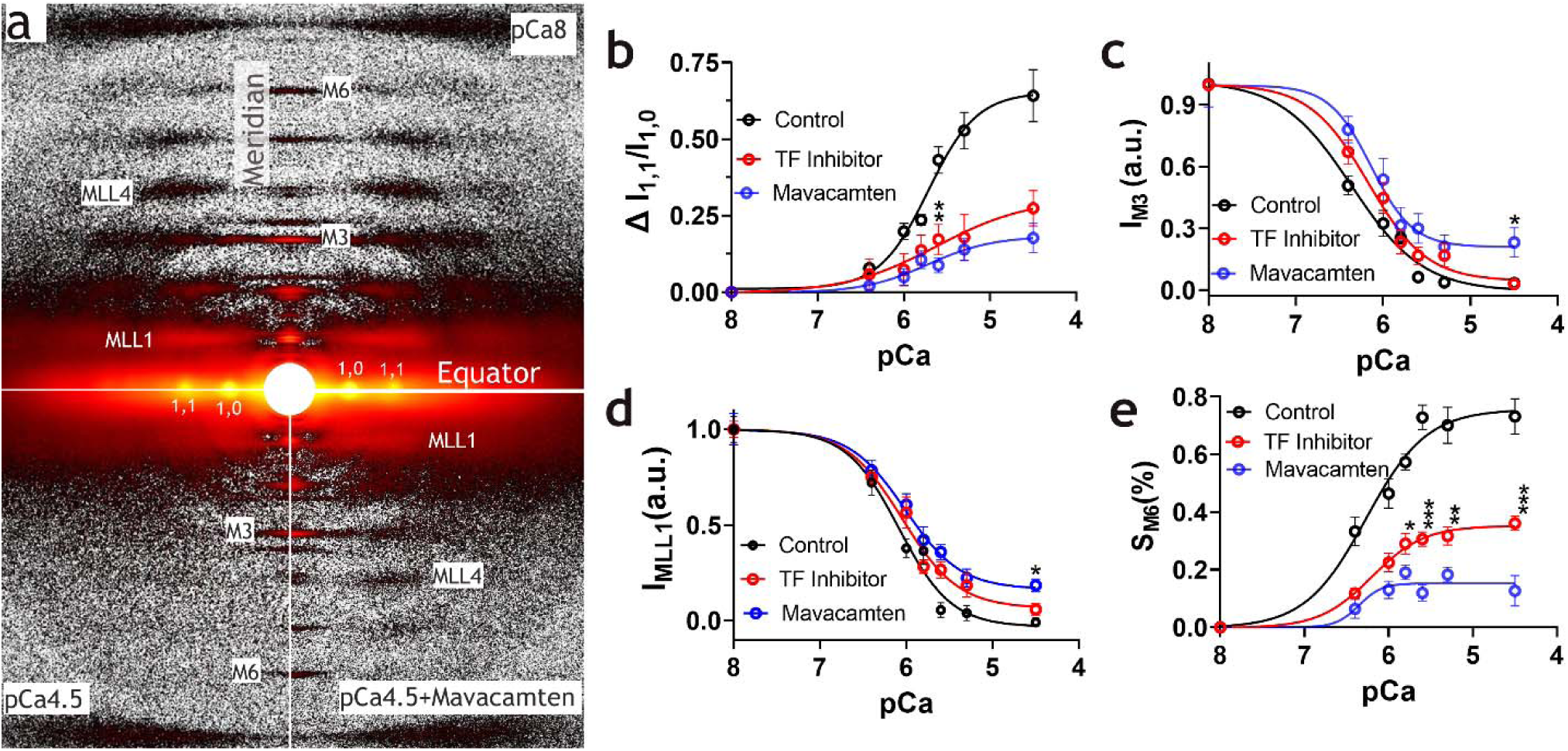
Thick filament structure changes in the presence of Ca^2+^. **a,** Representative X-ray diffraction patterns from permeabilized porcine myocardium under relaxing conditions (pCa 8, top panel) and activating conditions (pCa 4.5) with (bottom right) and without (bottom left) mavacamten in the absence of active force (treated with 100 μM MYK-7660). **b,** Change of equatorial intensity ratio (ΔI_1,1_/I_1,0_) at different Ca^2+^concentrations in the Control sub-group (black), thin filament (TF) inhibitor group (red), and the Mavacamten group (blue). The intensity of the third-order of myosin-based meridional reflection (**c**) and the intensity of the first-order myosin-based layer line (**d**) and in different concentrations of Ca^2+^ in the Control (black), TF inhibitor (red), and the Mavacamten (blue) groups. **e**, The spacing of the sixth-order myosin-based meridional reflection (S_M6_) in different Ca^2+^ concentrations in the Control (black), TF inhibitor (red), and the Mavacamten (blue) groups. The Ca^2+^ induced structural changes in the thick filament were blunted but not eliminated by the presence of mavacamten. * *p* <0.05, **: *p* <0.01, ***: *p* <0.001 between TF inhibitor and mavacamten groups. The results are given as mean ± SEM.

These Ca^2+^-mediated thick filament structural transitions in the presence of mavacamten, (blue curves in Fig. 2) have a pCa_50_ in the range of 5.9-6.4, essentially identical to the previously reported range in the absence of mavacamten (Control and TF Inhibitor group in Fig. 2). Consistent with the biochemical studies in Fig. 1, the amplitudes of the Ca^2+^-induced structural changes are the greatest in the Control group and the least in the Mavacamten group, while the TF inhibitor group is in between, indicating that even though mavacamten shifts more myosin heads into *off* states, these *off* state myosin heads are still sensitive to Ca^2+^.

### Mavacamten preserves positive force-frequency response in *vitro* and in *vivo*

As expected, and consistent with the Bowditch effect, increasing stimulation frequencies (from 1 to 4Hz) increases shortening fraction in isolated, membrane-intact, rat ventricular myocytes (Fig. 3a & 3b, Table S2, n=12); this Treppe effect was driven by increases in Ca^2+^ transient amplitudes (Fig. 3c, Table S2). Mavacamten (at 0.1 µM) preserved this Ca^2+^-mediated functional recruitment (from 3.9 ± 0.5% to 5.6 ± 0.3% ; (*p* = 0.0002)) despite markedly inhibiting shortening at each stimulation frequency. Indeed, comparable gains in function with pacing (i.e., linear slopes) were noted between control and mavacamten treated cells (+12 ± 3%/Hz vs. +13 ± 5%/Hz in VEH, n.s.). Higher stimulation frequencies also elevated diastolic cytosolic Ca^2+^ concentrations and resulted in progressive reductions in diastolic (resting) cell length (Fig. 3b, Table S2). Mavacamten preserved diastolic cell length (Fig. 3a), blunting the diastolic contracture (loss of diastolic length) with pacing (i.e., linear slope: -0.26 ± 0.04%/Hz (*p* = 0.00003) vs. -0.50 ± 0.07%/Hz in Control, (*p* = 0.00004); Fig. 3b) despite comparable increases in diastolic Ca^2+^-levels. This effect is likely mediated via the ability of mavacamten to facilitate mechanical re-lengthening (RT75: 86 ± 3%, or 0.15 ± 0.01s (*p* = 0.00002) vs. 0.17 ± 0.01s pre-dose (*p* = 0.00007)).

**Figure 3.**
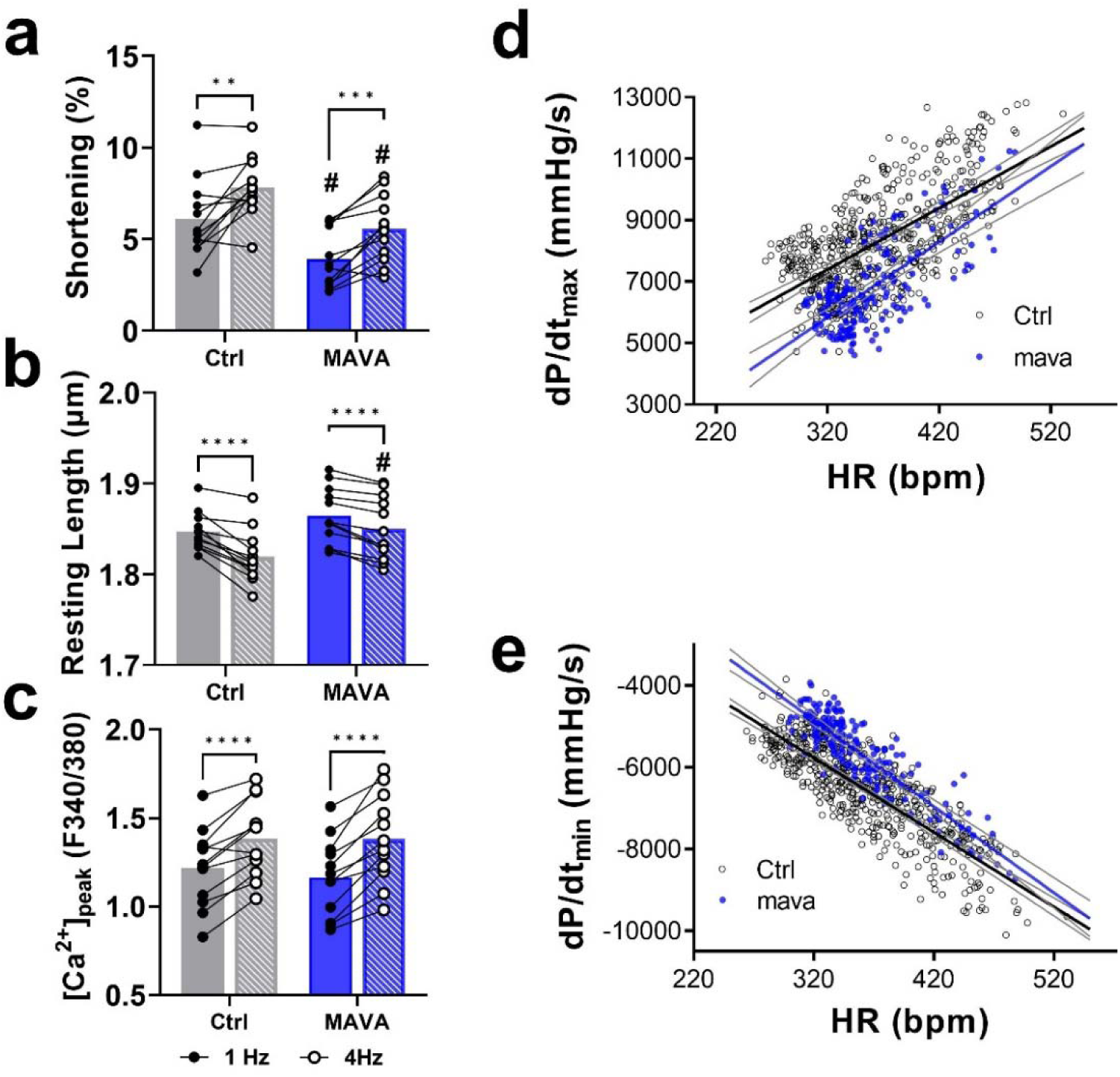
In *vitro* and in *vivo* Bowditch effect in the presence of mavacamten. In isolated rat ventricular myocytes, increases in stimulation frequency (from to 1 to 4 Hz) increasing shortening fraction (**a**) and decreasing resting length (**b**) by increase peak [Ca^2+^] during systole (**c**); In *vivo*, circadian changes in heart rate (HR) are accompanied by changes in the left-ventricular pressure derived dP/dt_max_ (**d**, direct) and dP/dt_min_ (**e**, inverse), reflecting HR-dependent function and mavacamten (1 mg/kg PO) preserved/enhanced these relationships. **: *p* <0.01, ***: *p* <0.001,****: *p* <0.0001. The bar graphs are given as mean values. **: *p* <0.01, ***: *p* <0.001,****: *p* <0.0001, # *p* <0.05 vs. Ctrl. Solid lines in panel **d** and **e** are simple linear regression fits and the dashed lines are 95% confidence bands of the best-fit line.

These in *vitro* observations were also noted *in vivo* in conscious, telemetered rats, where circadian changes in heart rate (over 24 h) were leveraged to establish the chronotropic-dependence of both dP/dt_max_ and dP/dt_min_, inotropic and lusitropic indices derived from the left-ventricular pressure (Fig. 3d). Under control conditions (N = 11), both a negative (strong) heart rate (HR)-dependence of dP/dt_min_ (-18.2 ± 0.5 mm Hg/s/bpm in Ctrl, R^2^=0.70, *p* < 0.0001) and a positive HR-dependence of dP/dt_max_ (20.0 ± 1.0 mm Hg/s/bpm in Ctrl, R^2^=0.41, *p* < 0.0001) were noted (black curve in Fig. 3e & 3f). Mavacamten administration (1 mg/kg PO, N = 6) preserved this HR-dependence (dP/dt_min_: -21.1 ± 0.9 mm Hg/s/bpm and dP/dt_max_: 24.6 ± 1.8 mm Hg/s/bpm_ notably, these relationships appeared steeper (*p* = 0.0077 vs. control slopes) under treatment. Taken together, these data demonstrates that mavacamten permits and/or enhances systolic and diastolic functional recruitment as HR increases, consistent with the Bowditch effect, but also with the Ca^2+-^dependent thick-filament recruitment described above.

### Stretch-mediated myosin filament structural changes are preserved in the presence of mavacamten

The systolic function of the heart is also regulated by its preload, determined in vivo by the degree of diastolic filling. At the cellular level, passive stretch of porcine myocardium can recruit reserve myosin heads from *off* states to *on* states promoting cross-bridge formation during activation (23). While it has been previously shown that mavacamten can stabilize the *off-state* of the thick filament and can blunt overall systolic gains with stretch (13, 23), it is unknown whether the stretch-mediated thick filament recruitment that mediates length-dependent activation remains operational in the presence of mavacamten. Here we demonstrate that both I_1,1_/I_1,0_ and the radius of the center of mass of the cross-bridges (R_m_), a direct measurement of the distance of helically ordered myosin heads from the thick filament backbone (52), increased significantly in both the control (N=2 and n=7; *p* < 0.05) and the mavacamten (N=2 and n=8; *p* < 0.05) group when muscle is stretched from a SL of 2.1 µm to 2.3 µm (Fig. 4 a & b, Table S3). Correspondingly, both I_MLL1_ and I_M3_ decreased significantly in both the control (N=2 and n=7; *p* < 0.05) and the mavacamten (N=2 and n=8; *p* < 0.05) groups when muscle is stretched from SL of 2.1 µm to 2.3 µm (Fig. 4 c & d, Table S3). The I_1,1_/I_1,0_ and R_m_ data indicate that myosin heads move radially away from the thick filament backbone concomitantly with a loss of helical ordering, indicated by I_MLL1_ and I_M3_, upon lengthening in the presence and absence of mavacamten. Remarkably, the SL lengthening induced changes are not significantly different (*p* > 0.05, Fig. 4, Table S3) between the control and mavacamten group strongly indicating that length-mediated structural changes in the myosin filaments are preserved in the presence of mavacamten.

**Figure 4.**
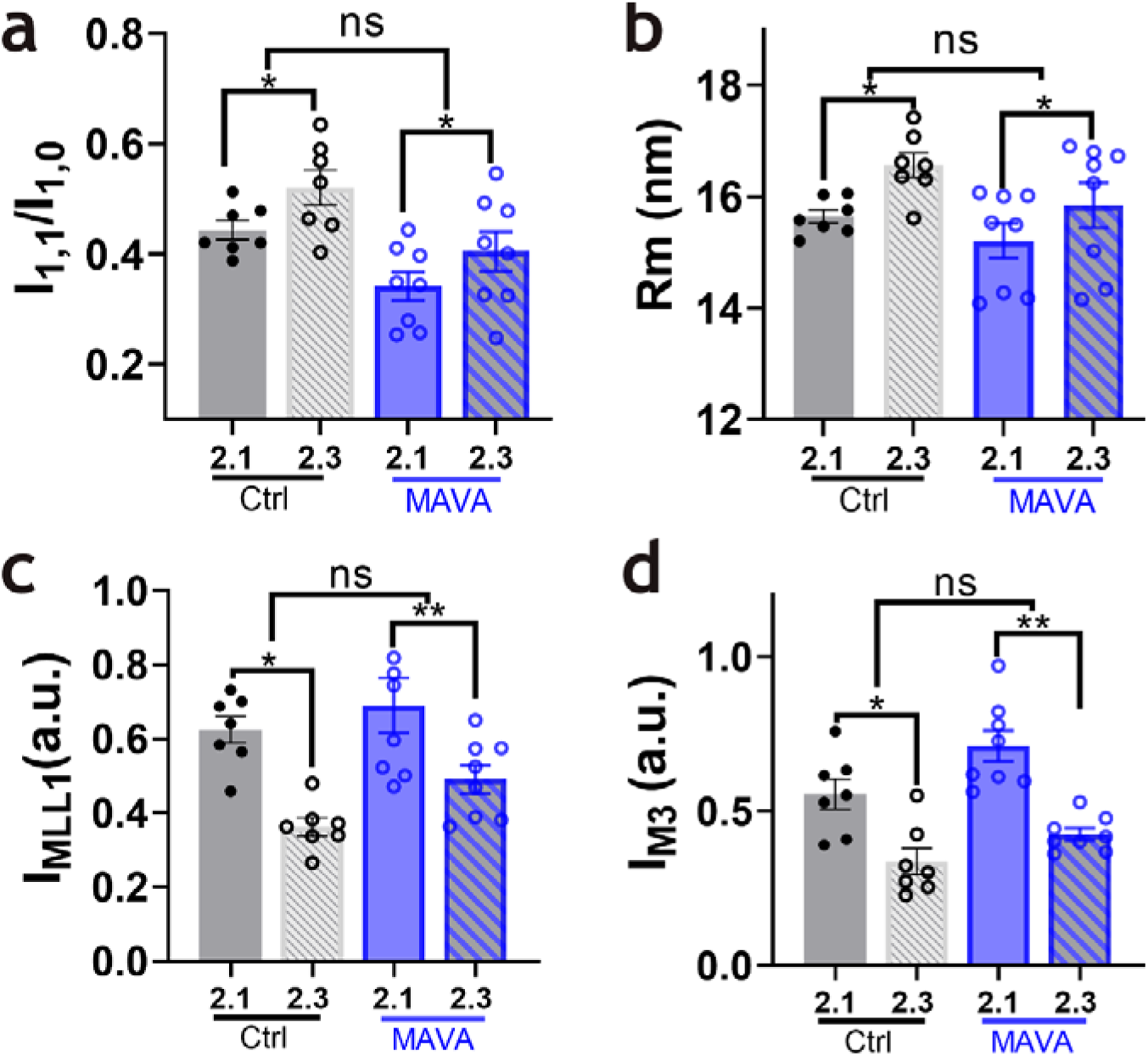
Length-dependent thick filament structural changes in the presence and absence of mavacamten. I_1,1_/I_1,0_ **(a)** and the radius of the average mass of myosin heads (R_m_) (**b)** increases while the intensity of the first-order myosin-based layer line (I_MLL1_) **(c)** and the intensity of the third-order myosin-based meridional reflection (I_M3_) **(d**) decreases at longer sarcomere length indicate structural transitions of the thick filament towards the *on* states upon sarcomere lengthening in the presence (blue) and absence (grey) of mavacamten. *: *p* <0.05, **: *p* <0.01. The results are given as mean ± SEM.

### β-adrenergic recruitment is preserved in the presence of mavacamten in *vivo*

In addition, the effects of β-adrenergic stimulation in the presence of mavacamten were studied *in vivo*. As expected at the supra-therapeutic dose-level used (ie, producing functional depression consistent with clinical finding at plasma concentrations >1000ng/mL, exceeding the range shown to be efficacious(14)), mavacamten resulted in marked functional inhibition in healthy pigs (N = 4), decreasing left ventricular ejection fraction (LVEF) by 32 ± 6% (from 72 ± 5 to 40 ± 5%, *p* = 0.0437, Fig. 5a), resulting in increased end-systolic and end-diastolic ventricular dimensions.. Mavacamten triggered significant increases in end-diastolic volume index (EDVI) from 41 ± 6 mL/m^2^ to 59 ± 6 mL/m^2^ (*p* =0.0263, Fig. 5b), which has been shown(27) to reflect increased ventricular distensibilitydue to its reduction in the number of residual diastolic cross-bridges. In this setting, β-AR stimulation with dobutamine restored LVEF towards baseline values (65 ± 4%, vs. DOSE (+mavacamten), *p* = 0.0437, Fig. 5a), decreasing end-systolic volumes (40 ± 3 to 24 ± 3 mL, *p* = 0.0302, Fig. 5c) and increasing stroke-volume (27 ± 5 to 45 ± 8 mL, *p* = 0.0471, Fig. 5d), suggesting the preservation of β-AR cardiac reserve. Notably, β-AR stimulation failed to decrease diastolic volumes (59 ± 6 % to 59 ± 6%, n.s.), suggesting that the diastolic effects of mavacamten remain unaltered. Taken together, these observations indicate that despite mavacamten-mediated thick-filament stabilization, a population of myosin-heads remains available to be recruited via β-adrenergic receptor stimulation.

**Figure 5.**
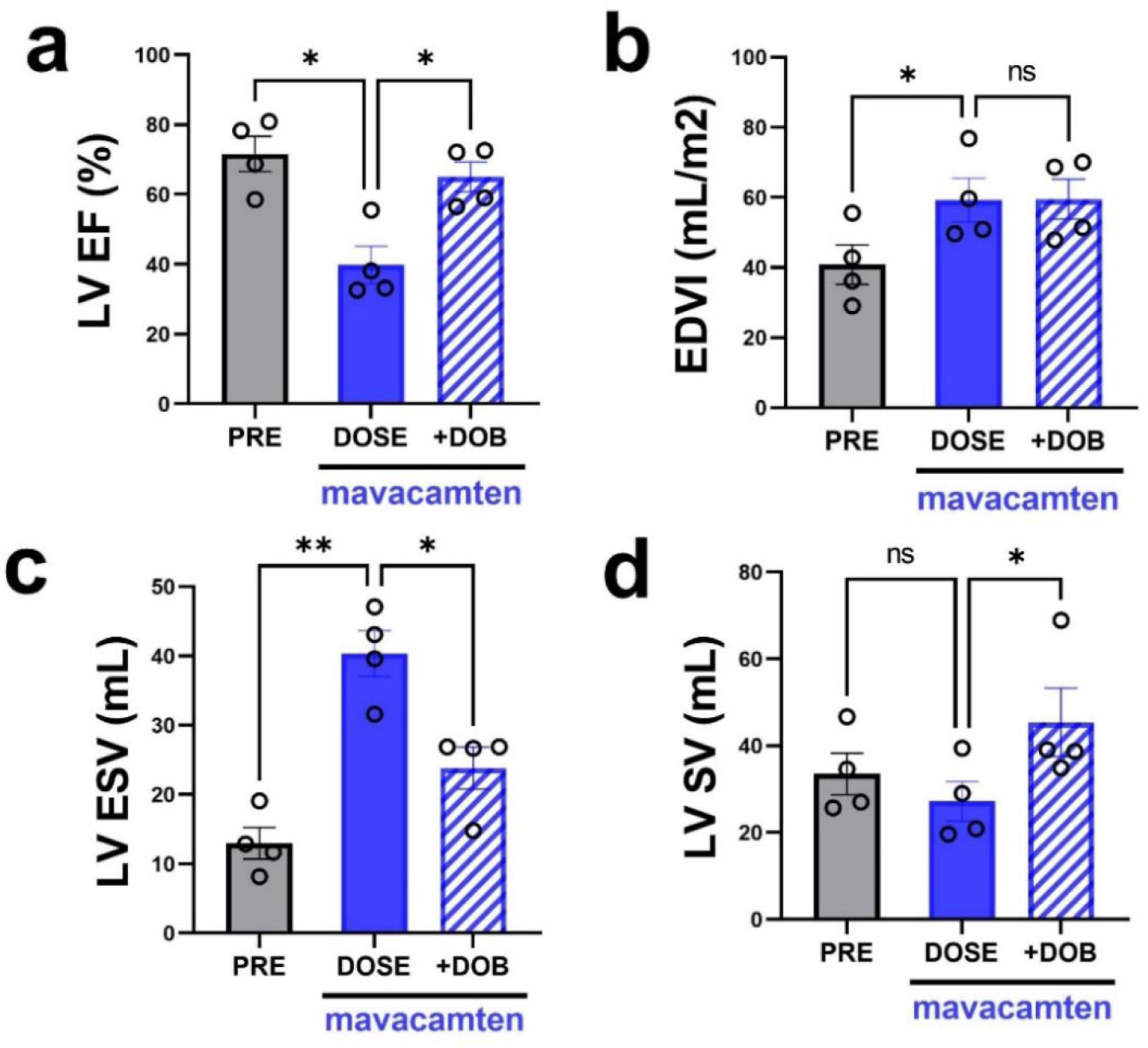
In *vivo* cardiac effects of mavacamten, and overlapping dobutamine, in healthy pigs. **a,** mavacamten (DOSE) at supra-therapeutic doses results in a marked decrease in left ventricular ejection fraction (LVEF) compared to PRE (no mavacamten), which are acutely recruitable by β-adrenergic receptor stimulation with dobutamine (+DOB; 5 µg/kg/min for 5 min). **b,** end-diastolic volumes index (EDVI) is increased by mavacamten and it is insensitive to β-AR stimulation. **c,** mavacamten increases LV end-systolic volume (LVESV) but β-AR stimulation is able to reverse it. **d**, mavacamten does not significantly affect LV stroke volume (LVSV) but β-AR stimulation significantly increases LVSV. ns: not significant, *: *p* <0.05, **: *p* <0.01. The results are given as mean ± SEM.

### Simulations of thick filament regulation by calcium

We used the computational simulation program MUSICO (see methods) to explore the effect of Ca^2+^ concentration on transitions in the population of myosin heads from sequestered *off* states to *on* states (47, 53). Simulations considered these transitions in the context of biochemical transitions from a "parked state" (PS) to the detached Myosin-ADP-phosphate (M.D.Pi) state and then allowing myosin to bind to actin to form the attached Actin-Myosin-ADP-phosphate (A.M.D.Pi) state. (Fig. 6). The “parked state” is functionally defined as any heads unable to bind to actin and would include any heads in the biochemically defined SRX state or the structurally defined *off* states regardless of whether these are the same or different populations. From the fraction of the myosin heads in the PS estimated from the X-ray diffraction data, assumed here to be proportional to the square root of the number of ordered heads, we establish the Ca^2+^-dependent state transition rate from the PS using the data from experiments involving the thin filament inhibitor MYK-7660 (thin filament inhibitor: Fig. 6, red symbols). We achieved very good matches to the data (Fig. 6; red line) by setting the rate of Ca^2+^ binding to very small values, i.e. mimicking the inhibition of opening thin filament regulatory units in the presence of MYK-7660, and setting parameters for the Ca^2+^-dependent state transition from the PS to M.D.Pi, *k_+PS_* (*Ca*^2+^), as: 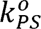 = 27 s^-1^, 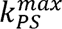 = 220 s^-1^ and Hill coefficient, *b* = 1.25. The reverse rate constant from M.D.Pi to PS is taken to be *k*_−*PS*_ = 200 s^-1^. Using parameters for the cross-bridge cycle slightly modified from simulations for human myocardium (46) and adjusted for temperature, species, and muscle preparation (i.e. demembranated vs. intact muscles, following (53)) the predicted changes in the fraction of the myosin heads in the PS as a function of Ca^2+^ concentration (pCa) (Fig. 6; black line) match the observed values for *off* state heads estimated from the X-ray diffraction experiments (Fig. 6; black symbols). We also explored the effect of 2 μM mavacamten on permeabilized porcine muscle in the absence of actin-bound cross-bridges (achieved with the thin filament inhibitor MYK-7660). The predicted fraction of myosin heads in the PS (Fig. 6, blue line) can be made to match the observations (Fig. 6, blue symbols), by changing only one rate constant k_−_*_PS_* from 200 to 263 s^-1^. This increase in *k*_−_*_PS_* reflects the activity of mavacamten in increasing the population of myosin heads in the PS at all Ca^2+^ concentrations.

**Figure 6.**
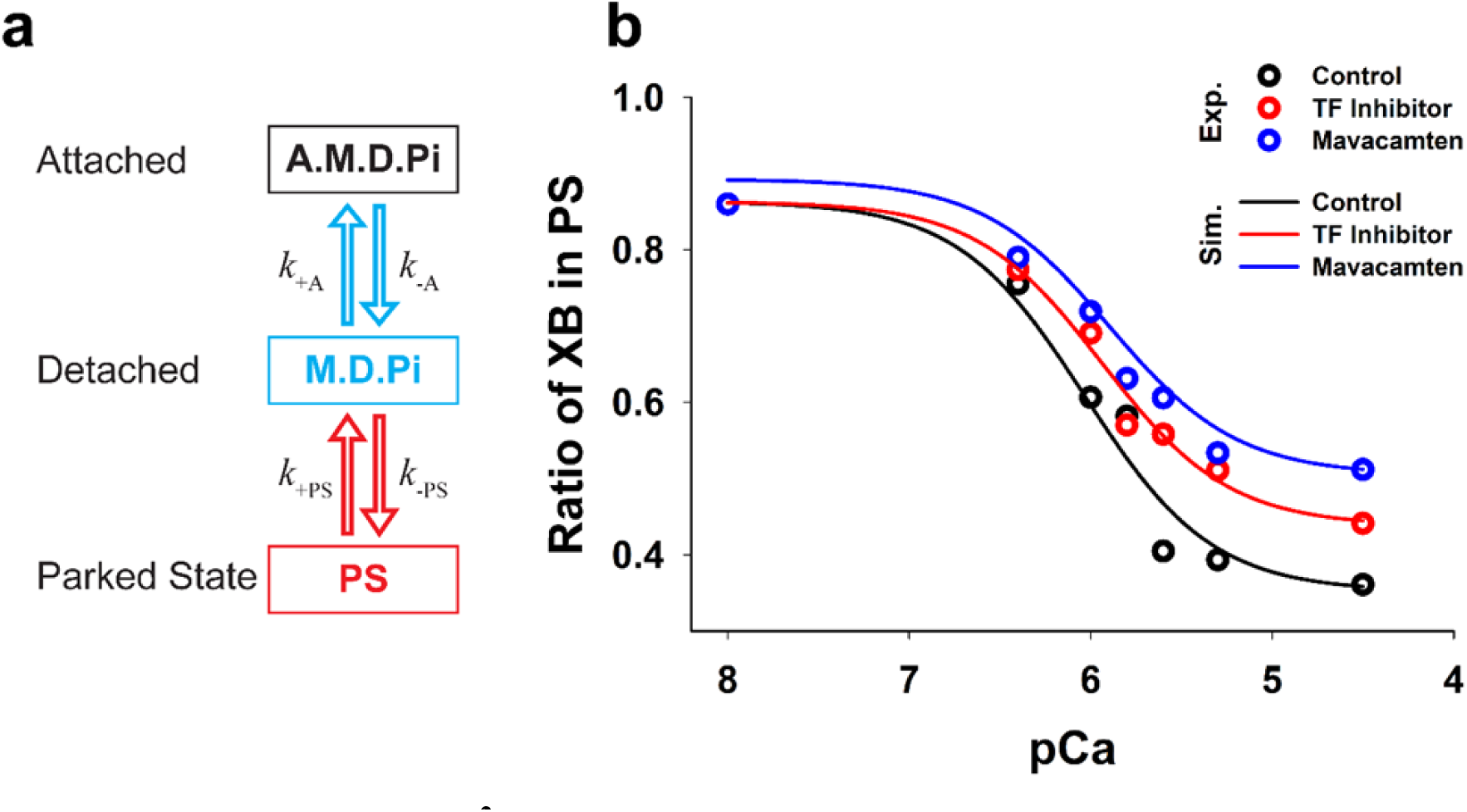
Simulation of the Ca^2+^-dependent parked state. **a,** Parked state (PS) myosin heads that are unavailable to form cross-bridges can be recruited by calcium to detached state(s) that are ready to participate in contraction. **b**, The population of PS heads decreases as the calcium concentration increases in both control muscle where active force is present (Exp.: experimental data; Sim.: simulation data) and thin filament inhibitor (TF inhibitor) group where active force i totally inhibited. In muscle where 2μM mavacamten is present, the population of the PS myosin heads decreases with increasing Ca^2+^ concentration, in the same manner as the control and TF inhibitor group, albeit with a reduced magnitude.

## Discussion

At the molecular level, excessive cross-bridge cycling due to mutations in sarcomeric proteins have been widely recognized as the primary contributor to hypercontractility in HCM(9, 10). Apart from the mutation-induced primary effect on myofilament function, other pathophysiological insults such as high ADP:ATP ratio or increased Ca^2+^ may add to the hypercontractile phenotype in HCM. Consistent with this hypothesis we have shown, in a recent report, that independent of thin filament-mediated activation, Ca^2+^ can directly destabilize the parked *off* states of myosin (24), thereby increasing the probability of excess actin-myosin interactions that could drive hypercontractility in HCM pathology, but also suggesting a mechanistic link between thick-filament-based myosin recruitment and inotropism.

Mavacamten is a small-molecule, FDA-approved, therapeutic for HCM that directly addresses this causal biology by directly targeting and inhibiting cardiac myosin (12) thus correcting the hypercontractility seen in HCM which can potentially prevent HCM associated symptoms and cardiac remodeling. It does so by sequestering myosin heads from the functionally more active DRX state to a less active SRX state, presumed to be structurally similar to the closed IHM-like state (13, 21, 25). Myosin heads in this state are thought to lie close to the thick filament backbones where they are quasi-helically ordered (the structurally defined *off* state) and away from the thin filaments so that they are unavailable to interact with actin (10, 54). Based on these two major propositions, a bigger question emerges. Does mavacamten permanently park these myosin heads in the *off* state(s)? Can these heads be recruited as and when required to boost the output of the heart? The answer(s) to these questions may provide a mechanistic explanation for the therapeutic benefits of mavacamten beyond mere obstruction relief, especially in the context of increased cardiac demand in patients during exercise.

This report shows that, at the molecular level, mavacamten shifts the SRX/DRX equilibrium towards the SRX state(s) and mitigates the deviation caused by the liberation of SRX heads to the DRX state(s) seen in HCM as indicated by the STF SRX assays. Our computational simulations predict the population of the myosin heads in the parked state (PS) in the presence of mavacamten, and a reasonable fit is achieved by changing only one rate constant *k*_−*PS*_ from 200 to 263 s^-1^. This result indicates that mavacamten promotes the transition of more myosin heads in the detached state into the sequestered PS. However, our simulation studies show that binding of mavacamten to myosin heads does not affect *k*_+*PS*_, suggesting that Ca^2+^ dependent recruitment of myosin heads from the PS is preserved in the presence of mavacamten. These results also serve to support the *in vitro* (Fig. 3 a & b) and *in vivo* observations (Fig. 3 d, e and f) that mavacamten preserves and/or enhances the Bowditch effect, as stimulation-frequency and heart-rate dependent recruitment of function was evident in the presence of mavacamten. It should also be noted that Matsubara and colleagues showed that the Bowditch, or staircase phenomenon in cardiac muscle is mediated by a frequency-dependent increase in the number of myosin projections transferred to the vicinity of the thin filaments during diastole (55) and during systolic contraction(56). Our studies provide a molecular mechanism for these findings, by associating the increases in myosin-mass transfer during Treppe with direct Ca^2+^-dependent regulation of *off* to *on* transitions of myosin on the thick filament.

We also noticed that while the change in SRX population was greater in mavacamten-treated STF compared to the control as Ca^2+^ concentration increases (Fig. 1b), the change in basal ATPase activity was actually larger for the control compared to the mavacamten group (Fig. 1c). It has been shown (22, 57) that mavacamten not only changes the SRX-DRX equilibrium to favor the SRX states but also slows down the ATPase rate of the DRX states of myosin. So, if A and k denote the normalized population and the ATPase rates of the different states of myosin, steady-state myosin ATPase is the weighted average defined as = A_SRX*_k_SRX_+A_DRX*_k_DRX_, where A_SRX_+A_DRX_= 1. Since mavacamten changes A_DRX_ and k_DRX,_ one can expect a more marked change in the steady-state ATPase measurements (as measured in Fig. 1c and 1d) as opposed to A_SRX_ (as measured in Fig. 1b).

In addition, at the level of the sarcomeres, both Ca^2+^ and passive lengthening can shift myosin heads from ordered *off* states close to the thick filament backbone to disordered *on* states closer to the thin filaments in the presence of saturating mavacamten (Fig. 4). Previous mechanical studies showed that mavacamten blunts but does not eliminate the systolic gains of LDA (23, 28, 29) supporting the notion that the myosin heads sequestered in the *off* SRX state(s) induced by mavacamten are available for length dependent recruitment. All these lines of evidence are supported by the observations from *in vivo* studies showing that mavacamten blunts but does not eliminate β-adrenergic stimulation effects on the heart.

Our data not only demonstrate that myosins in mavacamten induced *off* states in the thick filament are recruitable, thus leading to the preservation of the cardiac reserve, but also indicate that mavacamten, beyond mere LV outflow tract obstruction removal in oHCM patients, preserves cardiac-output recruitment mechanism(s) that enhance cardiac output in HCM patients. This hypothesis is supported by the observations in the phase II MAVERICK-HCM trial in patients without obstruction in which a high-risk subgroup with elevated myocardial injury biomarkers (cardiac troponin I >99^th^ percentile) or elevated diastolic filling pressures (average E/e’>14 on echocardiogram), one-third of mavacamten-treated patients (N =7 compared to zero in the placebo group, p = 0.03) met the composite functional endpoint—defined as achieving: 1) an improvement of at least 1.5 ml/kg/min in pVO_2_ with a reduction of ≥NYHA functional class; or 2) an improvement of 3.0 ml/kg/min or more in pVO_2_ with no worsening in NYHA functional class (18).

The ultimate function of the heart is to maintain adequate blood supplies under various conditions, especially during increased demand. Here we provide direct evidence, ranging from in *silico*, to in *vitro* and in *vivo*, that Ca^2+^-mediated activation, chronotropic inotropism, sarcomere stretch-mediated activation, and β-adrenergic receptor cardiac recruitment all remain active under mavacamten treatment, resulting in a likely maintenance (or in disease, enhancement) of cardiac reserve. These results not only provide a possible mechanistic explanation for the clinically observed increase in peak oxygen uptake (pVO_2_) with exercise in patients with HCM receiving mavacamten, but also lay the groundwork for a potential methodology to investigate the sarcomere basis of chronotropic incompetence in disease states motivate new therapeutic interventions.

### Limitations and future scope

Our current studies are limited to the short term recruitability of mavacamten stabilized myosin in healthy systems. We did not investigate how things might be in diseased myocardium. In clinical trials where HCM patients were administered mavacamten showed improved exercise capability beyond pVO_2_ strongly suggesting that the myosin stabilized by mavacamten is recruitable to accommodate increased physiological demands during exercise (58). It is worth noting that Ca^2+^-mediated activation, Bowditch effect, length-dependent activation, and beta-adrenergic stimulation are not separate phenomenon rather they have overlapping, complex mechanisms. For example, it has been shown that β-adrenergic stimulation has multiple effects: 1) it prolongs L-type calcium channel opening, effectively raising the Ca^2+^ concentration (59); 2) it phosphorylates multiple sarcomeric proteins including myosin-binding protein C (60), troponin I and C(61), and titin (62) all of which could modulate heart contractility. We also did not investigate the recruitability of mavacamten-stabilized myosin in the presence of other more commonly used drugs for HCM such as β blockers and calcium blockers (58), however, these agents operate at the sarcolemmal level and the effects shown here implicate direct actions of cytosolic Ca^2+^ and/or stretch on sarcomere proteins. It has been shown that pVO_2_ change was blunted in HCM patients on β blockers compared with patients that were not in the EXPLORER-HCM trial (58). We have shown that β-AR blockade does not alter the diastolic and systolic effects of mavacamten (63), The observations of this study, indicating that Ca^2+^ exerts are direct effect on the sarcomere, priming myosin heads for contraction provides a potential explanation for this clinical dichotomy, which could be attributed to the effects of β blockers on cardiac recruitment (64). All of these questions, while beyond the scope of current study, should be examined experimentally or clinically in the future.

## Acknowledgments

We thank Robert S. McDowell of Bristol Myers Squibb (formerly MyoKardia Inc.) for providing helpful comments to the manuscript. This research used resources of the Advanced Photon Source, a U.S. Department of Energy (DOE) Office of Science User Facility operated for the DOE Office of Science by Argonne National Laboratory under Contract No. DE-AC02- 06CH11357. This project is supported by grant P30 GM138395 from the National Institute of General Medical Sciences of the National Institutes of Health and the Ministry of Education, Science and Technological Development of the Republic of Serbia through Contracts No. 451- 03-68/2022-14/200378. The content is solely the authors’ responsibility and does not necessarily reflect the official views of the National Institute of General Medical Sciences or the National Institutes of Health.

## Author contributions

W.M, C.dR, S.N designed the experiments; W.M, H.G, C.dR, C.Z, R.S, S.G, S.N performed the experiments; W.M, L.Q, C.dD, C.Z. R.S, S.G, S.N. analyzed the data; M.P, S.M.M performed the simulations and interpreted the simulation results; W.M, S.M.M, C.dR, S.N, T.C.I. wrote the manuscript. All authors approved the final version of the manuscript.

## Data Availability

The dataset generated or analyzed during this study are included in this article. The raw data are available from the corresponding authors (Suman Nag; Carlos L. del Rio; Weikang Ma) on reasonable request.

## Code Availability

X-ray datasets were analyzed using data reduction programs belonging to the open-source MuscleX software package developed at BioCAT. Source codes are deposited on Github (https://github.com/biocatiit/musclex).

