## Supplementary material for "Myosin in autoinhibited *off* state(s), stabilized by mavacamten, can be recruited via inotropic effectors": SI

\# authors contributed equally

**Short title: mavacamten-stabilized myosins are recruitable**

* Corresponding authors:

Suman Nag,, 3160 Porter Drive Palo Alto, CA 94304;

Carlos L. del Rio,, 200 Technology Square Cambridge, MA 02139;

Weikang Ma,, 9700 S Cass Ave Argonne, IL 60439

**Supplemental Methods**

**Table S1. MUSICO parameters for the simulations of isometric contraction in skinned porcine muscles for the control case.**

| **Description** | **Parameter** | **Value** |
| --- | --- | --- |
| **Crossbridge Cycle** |  |  |
| Myosin-actin binding rate | $k_{+A}^{0}$ | 20 s^-1^ |
| Myosin-actin detachment rate^a^ | $k_{-A}^{0}$ | 40 s^-1^ |
| Myosin stroke forward cap rate constant | $k_{+Pi}^{cap}$ | 1000 s^-1^ |
| Myosin stroke reverse cap rate constant^b^ | $k_{-Pi}^{cap}$ | 10 s^-1^ |
| Power stroke Gibbs energy change ^1-3^ | $\Delta G_{stroke}$ | -11.3 $k_{B}T$ |
| Working stroke ^1-4^ | $d$ | 10.5 nm |
| Second working stroke ^1-3^ | $\delta$ | 1 nm |
| ADP release rate | $k_{+D}^{0}$ | 10 s^-1^ |
| ATP binding and myosin detachment rate constant^a^ | $k_{+T}$ | 10^6^ s^-1^ |
| Hydrolysis forward rate^a^ | $k_{+H}$ | 30 s^-1^ |
| Hydrolysis backward rate^a^ | $k_{-H}$ | 5 s^-1^ |
| Crossbridge stiffness ^1-4^ | $\kappa$ | 1.3 pN/nm |
| $k_{B}T$ at 30 °C | $k_{B}T$ | 4.185 pN·nm |
| **Parked State** |  |  |
| Transition rate constant to “parked state” | $k_{-PS}$ | 200 s^-1^ |
| Baseline rate constant | $k_{PS}^{0}$ | 27 s^-1^ |
| Amplitude | $k_{PS}^{max}$ | 220 s^-1^ |
| Calcium Hill function slope | $b$ | 1.25 |
| Half activation point of the Hill function | $\left[ Ca \right]_{50}$ | 2 μM |
| **Calcium Kinetics** |  |  |
| Calcium binding to TnC equilib. rate constant ^5, 6^ | $\tilde{K}_{Ca}$ | 10^6^ M^-1^ |
| Calcium binding rate constant to TnC ^5, 6^ | $\tilde{k}_{Ca}$ | 7.54·10^7^ M^-1^·s^-1^ |
| Calcium dissociation rate constant from TnC ^7-9^ | $k_{-Ca}$ | 75.4 s^-1^ |
| TnI-actin equilibrium rate const. at high Ca^2+^ | $\lambda$ | 10 |
| TnI-actin backward rate const. | $\lambda_{-}$ | 375 s^-1^ |
| TnI-actin-Ca cooperativity coefficient ^5, 6^ | $\varepsilon_{o}$ | 0.01 |
| **CFC** |  |  |
| Tropomyosin pinning angle ^10^ | $\phi_{-}$ | -25° |
| Myosin Tm angular displacement ^10^ | $\phi_{+}$ | 10° |
| Angular standard deviation of free CFC ^11, 12^ | $\sigma_{0}$ | 29.7° |
| Persistence length of Tm-Tn confined chain ^12^ | $1/\xi$ | 50 nm |
| **Sarcomere** |  |  |
| Length of sarcomere | $SL$ | 2.2 μm |
| Reference length of actin filament ^13, 14^ | $L_{a}^{o}$ | 1.1 μm |
| Interfilament spacing at SL=2.2 μm^15^ | $d_{10}$ | 41.7 nm |
| Thin filament elastic modulus ^16, 17^ | ${AE}_{a}$ | 65 nN |
| Thick filament elastic modulus ^16^ | ${AE}_{m}$ | 132 nN |

^a^ Based on mouse and human α-myosin values in ^18, 19^, with corrections for temperature, ionic strength as documented in ^20^.

^b^ The power stroke rates ($k_{+Pi}$ and $k_{-Pi}$) are expected to be slower in β-isoforms for ~5 fold and this is achieved by reducing adapting $G_{stroke}$ and decreasing $k_{-Pi}^{cap}$ by factor ~3; For the same isoform the power stroke rate increase from humans to rats and to mice is accomplished by increase in $k_{-Pi}^{cap}$.

**Sensitivity Analysis and Parameter Adjustments.** Most of the parameters used in MUSICO simulations are adopted from a previous study^21^ after adjustments from those appropriate to human to porcine ventricular trabeculae (Table SI 1). Several parameters were adjusted to match the observations of the parked state populations in control, with the TF inhibitor, and after 2μM mavacamten. These parameters were obtained from the sensitivity analysis choosing the parameter values for the experimentally observed values of parked state populations (Figure S1). The parameters $k_{PS}^{o}$ and $k_{PS}^{max}$defining $k_{+PS}\left( Ca \right)$, were obtained for the observed values of the PS with TF inhibitor for low (pCa 8.0) and for high calcium concentration (pCa 4.5) (Figures S1 A and B respectively). It is important to notice that these parameters are at the asymptotic tails of the PS population curves and their adjusted values do not affect other parameters defining $k_{+PS}\left( Ca \right)$, i.e. the estimated values for $k_{PS}^{o}$ and $k_{PS}^{max}$ can be determined from the PS value estimated from the X-ray diffraction data (Figs. S1 A and B). These estimates were made assuming that 100% of the heads are in the PS after treatment by 50 µm mavacamten at pCa 8^22^. To match the observations of the effect of mavacamten, only one state transition rate needs to be adjusted, namely $k_{-PS}$, by matching the PS population value at pCa 4.5 (Figure S1C). Similarly, to match the control data only the rate of myosin binding, $k_{+A}$, is adjusted by matching the PS population value in the control curve at pCa 4.5.

**Robustness of Stochastic Simulations.** The simulations of stochastic processes could show large variation in calculated values if the sampled population is small, thus for statistical averaging they require large number of runs.^4^ In MUSICO simulations we use a large population of interacting elements so the variation in calculated values is small. For example, in stochastic simulations limited to a half sarcomere with 500 myosin and 1,000 actin filaments, this is comparable with the number of filaments in a cross-section of a typical myofibril, provides sufficient statistical averaging without running the simulation multiple times.^1, 3, 21, 23^ In simulations of parked state populations for a half sarcomere we used here in preliminary simulations 200 half thick and 400 thin filaments the variation PS fraction in steady state was < 0.25%. For final simulations we used 500 half thick filaments and 1000 thin filaments which showed even smaller variation (< 0.12%). These variations are up to two orders of magnitude smaller than the standard deviation in experimental data demonstrating the robustness of the simulations.

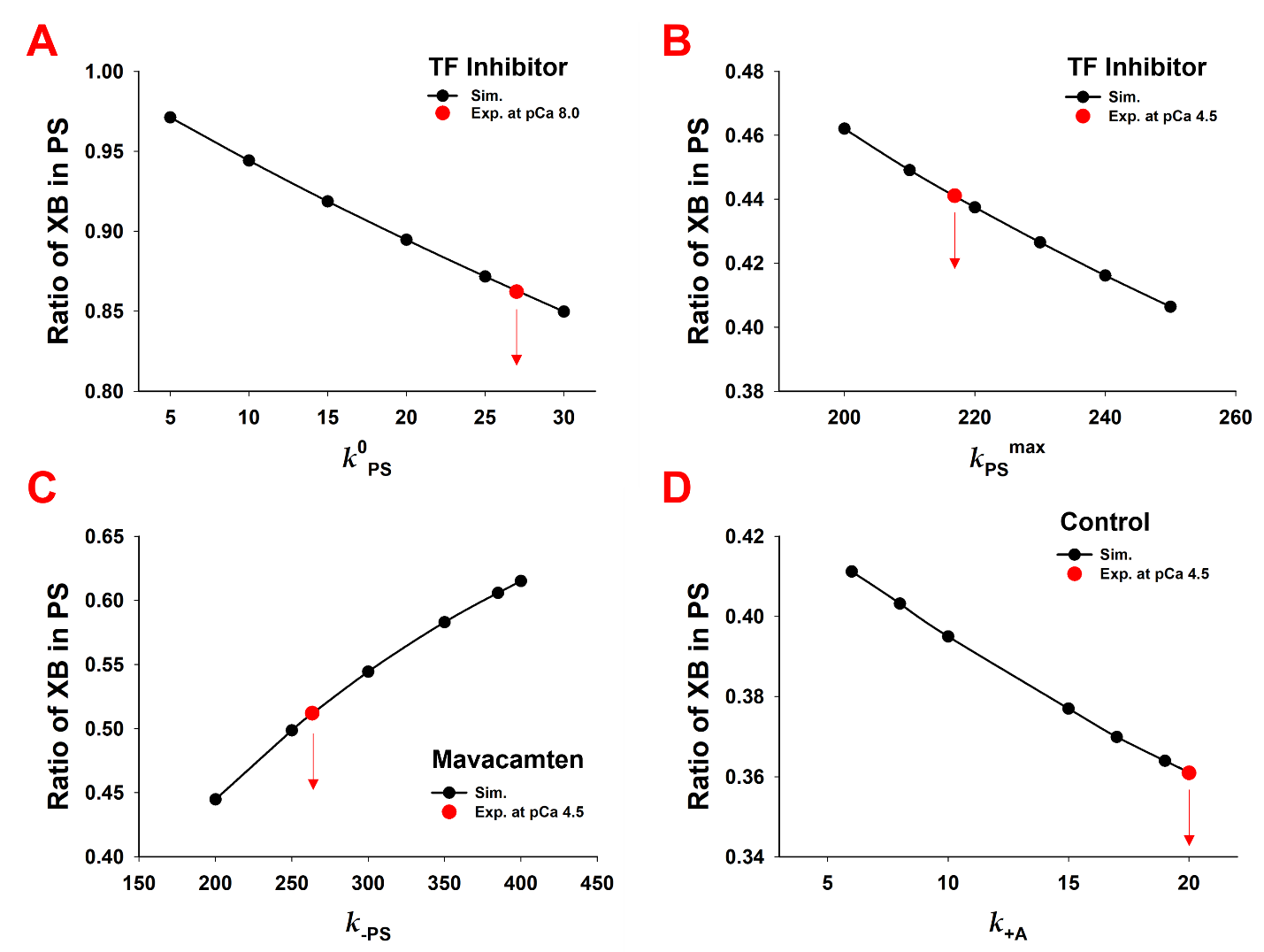

**Figure S1**. Sensitivity analysis of the fraction of crossbridges (XB) in PS on $k_{PS}^{o}$ , $k_{PS}^{max}$, $k_{-PS}$ and $k_{+A}$, where red circles represents the PS value estimated from the X-ray diffraction data; Estimated values of (A) $k_{PS}^{o}$ from the sensitivity relation of parked state population in presence of TF inhibitor at pCa 8.0; (B) Estimated value $k_{PS}^{max}$ in presence of TF inhibitor at pCa 4.5; (C) Estimated value $k_{-PS}$ in presence of TF inhibitor and 2 μM mavacamten at pCa 4.5; and (D) Estimated value of $k_{+A}$ from control at pCa 4.5.

**Supplemental Results**

**Table S2. Bowditch effect in the presence of mavacamten in rat ventricular myocytes**.

|  |  | **PRE** | **DOSE** | | |
| --- | --- | --- | --- | --- | --- |
|  |  | *@1Hz* | *@1Hz* | *@4Hz* | *Slope (@/Hz)* |
| SL (µm) | Ctrl | 1.72 ± 0.01 | 1.73 ± 0.01 | 1.68 ± 0.01^*^ | -0.50 ± 0.07 |
|  | MAVA | 1.74 ± 0.02 | 1.79 ± 0.02† | 1.75 ± 0.02 ^#,*^ | -0.26 ± 0.04^#^ |
| SF (%) | Ctrl | 6.6 ± 0.7 | 6.1 ± 0.6 | 7.8 ± 0.5^#,*^ | 13 ± 5 |
|  | MAVA | 6.6 ± 0.7 | 3.9 ± 0.5†,^#^ | 5.6 ± 0.5^#,*^ | 12 ± 3 |
| RT (ms) | Ctrl | 0.16 ± 0.01 | 0.15 ± 0.02 | 0.11 ± 0.01 | -9 ± 1 |
|  | MAVA | 0.17 ± 0.01 | 0.15 ± 0.02† | 0.11 ± 0.01 | -8 ± 1 |
| [Ca^2+^]peak (n/a) | Ctrl | 1.27 ± 0.06 | 1.22 ± 0.06 | 1.39 ± 0.06^*^ | 5.6 ± 0.6 |
|  | MAVA | 1.24 ± 0.06 | 1.17 ± 0.06 | 1.38 ± 0.07^*^ | 6.3 ± 0.9 |
| [Ca^2+^]min (n/a) | Ctrl | 0.74 ± 0.03 | 0.73 ± 0.03 | 0.85 ± 0.03^*^ | 6.4 ± 0.7 |
|  | MAVA | 0.71 ± 0.02 | 0.69 ± 0.02† | 0.84 ± 0.03^*^ | 6.9 ± 0.7 |

DOSE: cells randomly assigned to either mavacamten at 0.1µM (MAVA) or time-matched control (Ctrl) for 5min (n = 12 cells each). SL: resting (diastolic) cell length; SF: Shortening fraction; RT: relaxation time (to 75% of baseline). Peak and minimal (diastolic) Ca^2+^ concentrations derived from the F340/380 ratios.

†: P < 0.05 vs. PRE; #: P < 0.05 vs. Ctrl; and *: P < 0.05 vs. 1Hz (under DOSE)

**Table S3. Length-dependent thick filament structural changes in the presence and absence of mavacamten.**

| SL (μm) | 2.1 | 2.3 | *p* (2.1vs.2.3) | 2.3 - 2.1 | *p* (Ctrl vs. MAVA) |
| --- | --- | --- | --- | --- | --- |
| Ctrl_I_1,1_/I_1,0_ | 0.44 ± 0.017 | 0.52 ± 0.032 | *p* = 0.016 | 0.06 ± 0.015 | *p* = 0.87 |
| MAVA_I_1,1_/I_1,0_ | 0.34 ± 0.025 | 0.40 ± 0.035 | *p* = 0.016 | 0.08 ± 0.02 |  |
| Ctrl_R_m_ (nm) | 15.64 ± 0.12 | 16.56 ± 0.22 | *p* = 0.016 | 0.64 ± 0.17 | *p* = 0.46 |
| MAVA_R_m_ (nm) | 15.21 ± 0.31 | 15.84 ± 0.11 | *p* = 0.008 | 0.92 ± 0.20 |  |
| Ctrl_I_MLL1_ | 0.62 ± 0.04 | 0.36 ± 0.02 | *p* = 0.016 | -0.29 ± 0.04 | *p* = 0.40 |
| MAVA_I_MLL1_ | 0.69 ± 0.07 | 0.49 ± 0.04 | *p* = 0.008 | -0.22 ± 0.04 |  |
| Ctrl_I_M3­­_ | 0.56 ± 0.05 | 0.34 ± 0.04 | *p* = 0.016 | -0.20 ± 0.06 | *p* = 0.40 |
| MAVA_I_M3­­_ | 0.71 ± 0.05 | 0.42 ± 0.02 | *p* = 0.008 | -0.26 ± 0.04 |  |

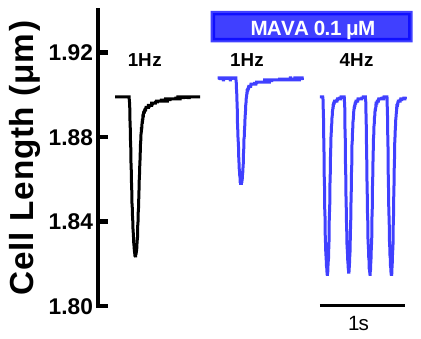

**Figure S2**. Representative tracings shows that mavacamten preserves frequency-dependent functional recruitment but blunts diastolic contracture
